## Supplementary Information for "Break-up and Recovery of Harmony between Direct and Indirect Pathways in The Basal Ganglia; Huntington’s Disease and Treatment"

**Sang-Yoon Kim · Woochang Lim**

**Abstract** This is the Supplementary Information (SI) for "Break-up and Recovery of Harmony between Direct and Indirect Pathways in The Basal Ganglia; Huntington's Disease and Treatment." In this SI, we briefly describe a spiking neural network of the basal ganglia, considered in our recent work (Kim and Lim, 2023).

### 1 Spiking Neural Network for The Basal Ganglia

Recently, based on the spiking neural networks for the basal ganglia (BG) developed in previous works (Humphries et al., 2009; Tomkins et al., 2014; Fountas and Shanahan, 2017), we made refinements on the BG spiking neural network to become satisfactory for our study to quantify harmony between direct and indirect pathways for the healthy and Parkinsonian states (Kim and Lim, 2023). Details on the BG spiking neural network are given in Sec. 2 and Appendices in (Kim and Lim, 2023). Here, we make brief description on the BG spiking neural network; for more details, refer to Sec. 2 and Appendices in (Kim and Lim, 2023).

Figure 1 shows a box diagram of major neurons and synaptic connections in the BG spiking neural network. Based on the anatomical property of the BG (Oorschot, 1996; Bar-Gad et al., 2003; Mailly et al., 2003; Sadek et al., 2007), we consider the BG spiking neural network, composed of D1/D2 spine projection neurons (SPNs), subthalamic nucleus (STN) neurons, globus pallidus (GP) neurons, and substantia nigra pars reticulata (SNr) neurons. For more details on the numbers of the BG cells and their synaptic connection probabilities, refer to Sec. IIA and Tables I and II in (Kim and Lim, 2023).

S.-Y. Kim
Institute for Computational Neuroscience and Department of Science Education, Daegu National University of Education, Daegu 42411, Korea


W. Lim (corresponding author)
Institute for Computational Neuroscience and Department of Science Education, Daegu National University of Education, Daegu 42411, Korea


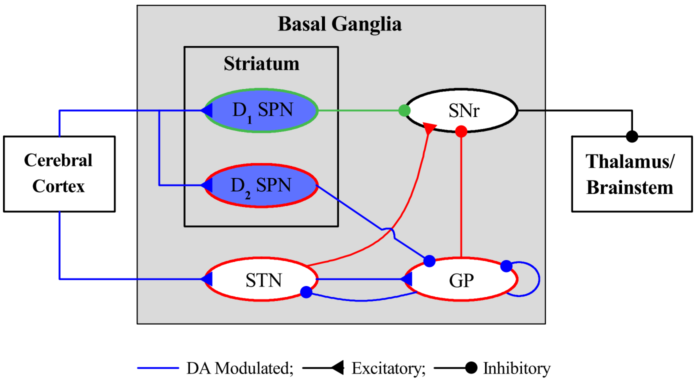


**Fig. 1** Box diagram of our spiking neural network for the basal ganglia (BG). Excitatory and inhibitory connections are denoted by lines with triangles and circles, respectively, and dopamine-modulated cells and connections are represented in blue color. Striatum and STN (subthalamic nucleus), receiving the excitatory cortical input, are two input nuclei to the BG. In the striatum, there are two kinds of inhibitory spine projection neurons (SPNs); SPNs with the D1 receptors (D1 SPNs) and SPNs with D2 receptors (D2 SPNs). The D1 SPNs make direct inhibitory projection to the output nuclei SNr (substantia nigra pars reticulata) through the direct pathway (DP; green color). In contrast, the D2 SPNs are connected to the SNr through the indirect pathway (IP; red color) crossing the GP (globus pallidus) and the STN. The inhibitory output from the SNr to the thalamus/brainstem is controlled through competition between the DP and IP.

Next, we make brief descriptions on the single neuron models and the dopamine (DA) effects in the BG spiking neural network; for details refer to Sec. IIB and Appendix A in (Kim and Lim, 2023). As the single neuron model, we use the Izhikevich spiking neuron model (which is not only biologically plausible, but also computationally efficient) as the elements of the BG spiking neural network (Izhikevich, 2003, 2004, 2007a,b). The BG spiking neural network consists of 5 populations of D1/D2 SPNs, the STN, the GP, and the SNr; for parameter values of each BG cells, refer to Table III in (Kim and Lim, 2023). The modulation effect of dopamine (DA) on the D1/D2 SPNs are also considered (Humphries et al., 2009; Tomkins et al., 2014; Fountas and Shanahan, 2017). For details, refer to Sec. IIB, Appendix A, and Table IV in (Kim and Lim, 2023).

The state of a neuron in each population is characterized by its membrane potential and slow recovery variable. Time-evolution of the membrane potential and the slow recovery variable is governed by 3 kinds of currents into the neuron such as the external current from the external background region, the synaptic current, and the injected stimulation current. Here, we consider the case of no injected stimulation current. The external current is modeled in terms of spontaneous (in-vivo) current (to get the spontaneous in-vivo firing rate) and random background input; for more details, refer to Sec. IIB, Appendix A, and Table V in (Kim and Lim, 2023).

Finally, we consider the synaptic currents and the DA effects; detailed explanations are given in Sec. IIB and Appendix B in (Kim and Lim, 2023). There are 3 kinds of synaptic currents from a presynaptic source population to a postsynaptic neuron in the target population; 2 kinds of excitatory AMPA and NMDA receptor-mediated synaptic currents and one type of inhibitory GABA receptor-mediated synaptic current. For each $R$ (AMPA, NMDA, and GABA) receptor-mediated synaptic current, the synaptic conductance is given by a product of the maximum synaptic conductance, the average number of afferent synapses, and the fraction of open postsynaptic ion channels. The time course of fraction of open ion channels is provided by a sum of exponential functions over presynaptic spikes. The synaptic parameters are given in Table VI in (Kim and Lim, 2023). These synaptic parameter values are based on physiological property (Park et al., 1982; Nakanishi et al., 1990; Fujimoto and Kita, 1993; Góngora-Alfaro et al., 1997; Götz et al., 1997; Richards et al., 1997; Bevan and Wilson, 1999; Bevan et al., 2000; Dayan and Abbott, 2001; Bevan et al., 2002; Liu et al., 2022; Hallworth et al., 2003; Baufreton et al., 2005; Wolf et al., 2005; Shen and Johnson, 2006; Moyer et al., 2007; Gertler et al., 2008; Bugaysen et al., 2010; Connelly et al., 2010; Ammari et al., 2011). The modulation effect of DA on afferent synapses into the D1/D2 SPNs, the STN, and the GP is also taken into consideration (Humphries et al., 2009; Tomkins et al., 2014; Fountas and Shanahan, 2017); for details, refer to Table VII in (Kim and Lim, 2023).
